# Neural Circuit Remodeling Underlying Enhanced Feeding During Pregnancy in Mice

**DOI:** 10.1101/2025.08.07.669199

**Authors:** Kengo Inada, Gen-ichi Tasaka, Mitsue Hagihara, Kenta Kobayashi, Hiroko Yukinaga, Kazunari Miyamichi

**Affiliations:** Laboratory for Comparative Connectomics, RIKEN Center for Biosystems Dynamics Research, Kobe, Hyogo 650-0047, Japan; Department of Viral Vector Development, National Institute for Physiological Sciences, Okazaki, Aichi 444-8585, Japan; CREST, Japan Science and Technology Agency, Kawaguchi, Saitama 332-0012, Japan; Graduate School of Science, University of Hyogo, Akou-gun, Hyogo 678-1297, Japan

## Abstract

Pregnancy represents a unique physiological state marked by substantial adaptations influencing both behavior and metabolism^1^. Alterations have been observed across multiple levels—from changes in cellular morphology, synaptic density, and neuroglial interactions to a brain-wide reduction in gray matter volume^2–7^. While these adaptations are thought to support offspring survival and promote maternal physiological and psychological well-being^8^, their functional significance remains largely unresolved. In particular, linking localized synaptic plasticity to a broader brain-wide circuit dynamics has posed a major challenge. Using virus-mediated screening in mice, we investigated the pregnancy-induced remodeling of the presynaptic landscape of paraventricular hypothalamic (PVH) oxytocin neurons, a central hub of maternal physiology^9,10^. Here we show a selective and reversible elimination of excitatory synaptic inputs from the medial preoptic nucleus to PVH oxytocin neurons, displaying region-, cell-type-, and pathway-specificity. This reduction in excitatory drive attenuated the anorexigenic activity of oxytocin neurons during feeding, thereby promoting the increased food intake characteristic of pregnancy. These data demonstrate that the selective suppression of anorexigenic PVH oxytocin neuron activity plays a critical role in appetite regulation during pregnancy. More broadly, our data identify a pregnancy-associated remodeling of hypothalamic neural circuits with direct physiological relevance.

---

The adult brain exhibits remarkable flexibility in adapting physiology and behavior to meet life-stage-specific demands. During pregnancy, for instance, food intake increases to support fetal development^1^, while maternal caregiving behaviors are facilitated in preparation for infant rearing^2^. These adaptations are thought to involve changes in synaptic strength. Recent seminal studies have demonstrated estrous cycle-dependent periodic remodeling of hypothalamic neural connectivity in female mice^11^ and social experience-dependent reorganization of limbic neural circuits in male mice^12^, underscoring the plasticity of these networks. However, how brain-wide neural circuit dynamics are shaped during pregnancy remains poorly understood. To address this knowledge gap, we utilized a virus-mediated monosynaptic tracing approach^13,14^ to map presynaptic inputs to oxytocin (OT) neurons in the paraventricular hypothalamus (PVH), which play key roles in parturition, lactation, maternal care, social bonding, and regulation of feeding and energy expenditure^9,10^.

## Remodeling of neural connections to PVH OT neurons during pregnancy

To explore the potential mechanisms regulating PVH OT neuron activity during pregnancy, we utilized rabies virus-based retrograde transsynaptic tracing^15^ of PVH OT neurons in mice^14^ (Fig. 1a). Starter cells, defined by the overlap of rabies virus-derived green fluorescent protein (GFP) and TVA-mCherry, were generated in age-matched non-pregnant (NP) females and females on gestation day (GD) 13. Because the rabies virus spreads within 1 or 2 days following infection^16^ and labeling patterns stabilize after 4 days^15^, this experimental scheme reflects the input landscape during late pregnancy, corresponding to GDs 15–17. Although the number of starter cells was comparable between these two conditions (Fig. 1b), we observed sparser GFP labeling in specific brain regions such as the medial preoptic nucleus (MPN) and anteroventral periventricular nucleus (AVPV) in pregnant compared to NP females (Fig. 1c). Conversely, labeling in other brain regions, such as the dorsomedial hypothalamus (DMH) and lateral hypothalamic area (LHA), remained unchanged. The convergence index, defined as the number of GFP-expressing presynaptic neurons in each brain region normalized to the number of starter cells, was significantly reduced in the MPN, whereas the LHA-derived input remained largely unchanged (Fig. 1d). These data suggested a region-specific reduction in the input of PVH OT neurons during late pregnancy.

**Fig. 1.**
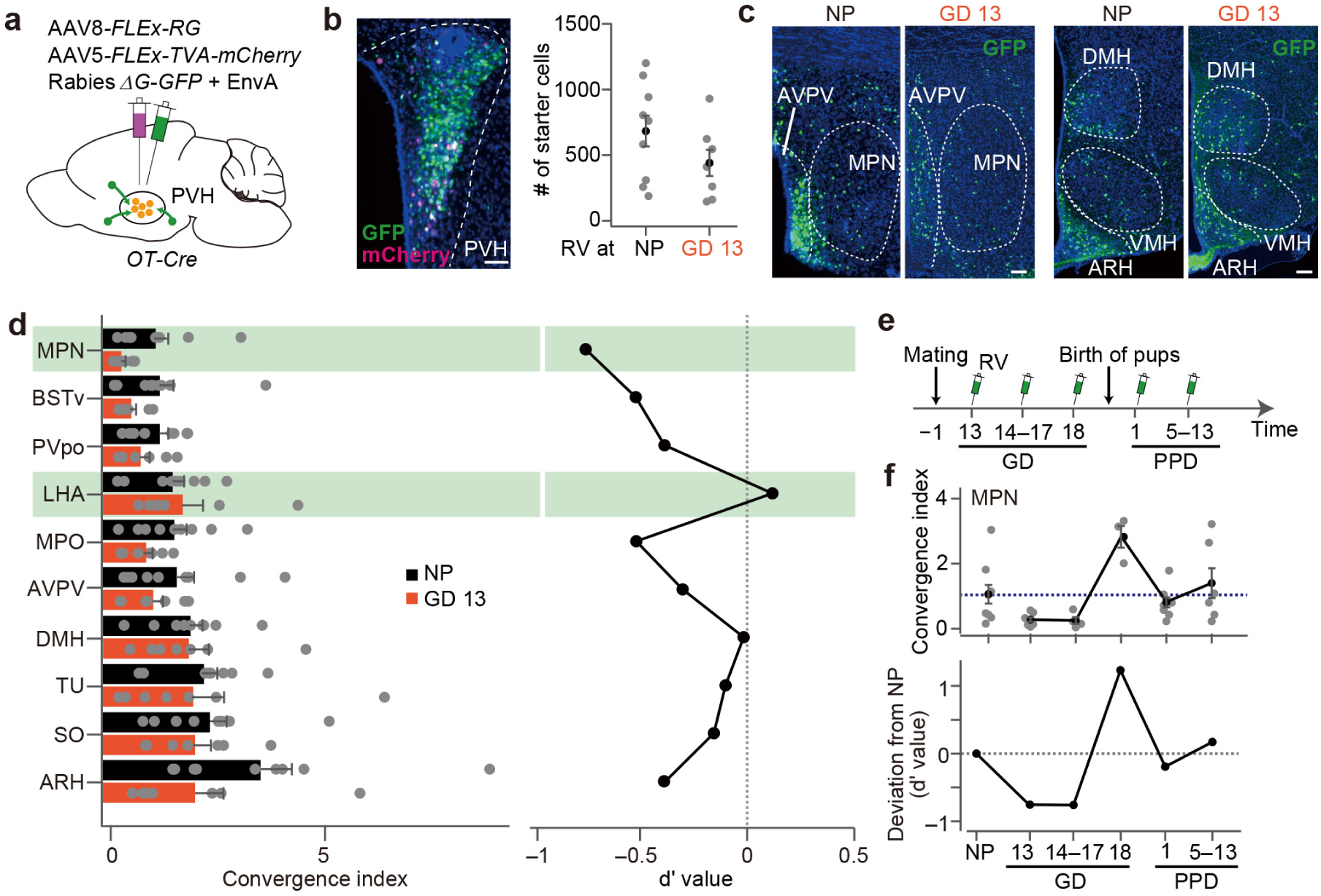
Dynamics of presynaptic landscape of PVH OT neurons during pregnancy. **a**, Schematic of the virus injections. Rabies virus was injected on gestation day (GD) 13. **b**, Left: Representative coronal section containing starter cells (white) defined as the overlap of mCherry (magenta) and GFP (green). Blue, DAPI. Scale bar, 50 μm. Right: Number of starter cells in the PVH in non-pregnant (NP) and pregnant groups (p > 0.17, two-tailed Student’s *t*-test). N = 9 and 7 mice for NP and pregnant groups, respectively. **c**, Representative coronal sections of presynaptic cells revealed by rabies-GFP. Scale bar, 100 μm. **d**, Normalized inputs from the top 10 nuclei to the PVH OT neurons in NP and pregnant females (left) and quantification of the difference in convergence index (right). A d′ value smaller than 0 represents a single PVH OT neuron in pregnant females receiving lesser input. The green shadows highlight MPN and LHA. **e**, Timeline of the time course analysis. The green syringe indicates the timing of rabies virus injection. PPD, postpartum day. **f**, Convergence index of MPN labeling (top) and d′ values calculated between each condition and NP (bottom). N = 4–9 mice per condition. The dotted line represents the average of NP females (from the same dataset shown in panel **d**). Error bars: standard error of the mean (SEM). See Table S1 for abbreviations and Tables S2 and S3 for the raw dataset. We note that 5 out of 9 NP mice data and all PPD 1 data were obtained from our previous publication^45^.

To assess the reversibility of this reduction, we conducted trans-synaptic tracing at multiple points throughout pregnancy and after parturition (Fig. 1e). Inputs originating from the MPN showed a transient decline in convergence index during late pregnancy, followed by a rapid resurgence in labelling around parturition, as revealed by rabies-virus injections on GD 18 (Fig. 1f). In the postpartum phase, MPN connectivity returned to values comparable to those of the NP group. These observations indicate that PVH OT circuits in the female brain undergo transient, region-specific remodelling during pregnancy.

To examine the electrophysiological properties, we performed targeted patch-clamp recordings of PVH OT neurons visualized via an AAV that drives mCherry in an OT neuron-specific manner^14,17^ in age-matched NP and GD 15 females. The input resistance measured at the soma and the membrane time constant exhibited no significant differences between the NP and GD 15 females (Extended Data Fig. 1a, 1b). Conversely, PVH OT neurons in GD 15 females demonstrated a significantly lower frequency of spontaneous excitatory postsynaptic currents (sEPSCs), whereas spontaneous inhibitory postsynaptic currents (sIPSCs) remained stable (Extended Data Fig 1c–e). Moreover, the PVH OT neurons in GD 15 females were more likely to receive sEPSCs of smaller amplitudes (Extended Data Fig. 1f), further supporting an overall reduction in excitatory drive.

Next, we performed channelrhodopsin 2 (ChR2)-assisted circuit mapping (CRACM) in acute brain slices^18^ to analyze synaptic transmission from the MPN. This region exhibited the most prominent reduction in rabies labeling (Fig. 1) in PVH OT neurons, with the LHA serving as the control region that showed no substantial change. To distinguish between cell types, we used *vGluT2-Cre* for excitatory neurons and *vGAT-Cre* for inhibitory neurons (Figs. 2a, Extended Data Fig. 1g). An AAV vector driving Cre-dependent ChR2 was selectively injected into the MPN (Fig. 2b, Extended Data Fig. 2). In *vGluT2-Cre* mice, a brief blue light pulse evoked a short-latency excitatory response in the PVH OT neurons (Fig. 2c), which was abolished by glutamate receptor antagonists (Extended Data Fig. 2e), confirming the monosynaptic glutamatergic transmission. In *vGAT-Cre* mice, the same optical stimulation elicited an inhibitory response in PVH OT neurons (Fig. 2d). The connection ratio from MPN *vGluT2*+ neurons to PVH OT neurons was significantly lower in GD 15 females. Furthermore, the amplitude of ChR2-evoked EPSCs in the connected PVH OT neurons was also reduced (Fig. 2c). In contrast, no differences were observed in the connection ratio or response amplitude of inhibitory transmissions (Fig. 2d).

**Fig. 2.**
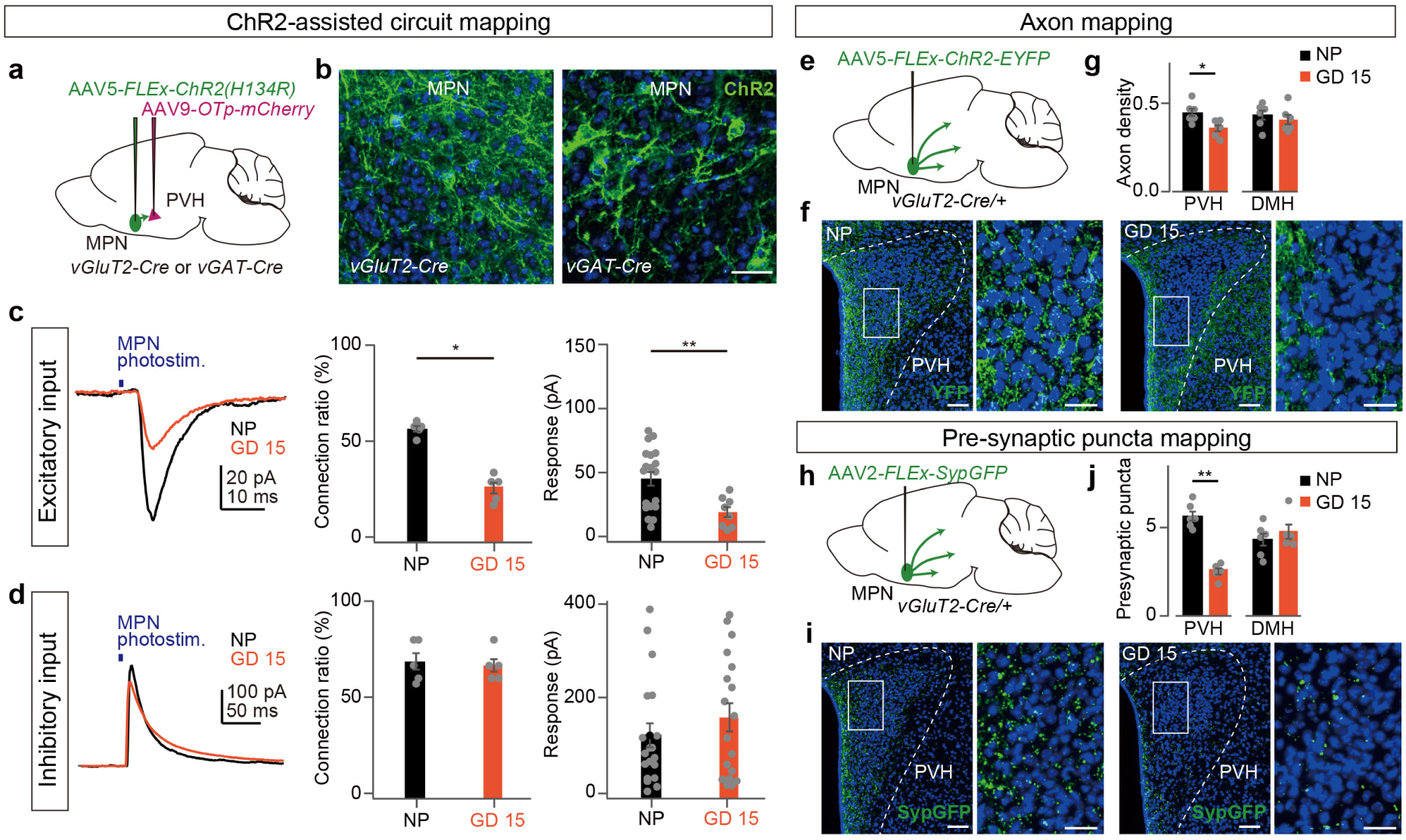
Excitatory synaptic remodeling in MPN-to-PVH OT neuron circuit during pregnancy. **a**, Schematic of the virus injections for ChR2-assisted circuit mapping. **b**, Representative coronal section of *vGluT2+* (left) or *vGAT+* (right) neurons in the MPN. Green is ChR2-EYFP amplified by anti-GFP staining. Blue, DAPI. Scale bar, 20 μm. **c**, **d**, Left: Representative responses from PVH OT neurons to optogenetic activation of *vGluT2*+ (**c**) or *vGAT+* (**d**) axons originating from MPN neurons. Each trace is an average of 10 trials. Blue bar, light stimulation (1 ms, 3.6 mW). Middle: The ratio showing that the recorded OT neuron was connected to the stimulated neurons. N = 5 mice each. *p < 0.05, two-sided Mann-Whitney *U*-test. Right: The light-induced response of PVH OT neurons. The excitatory response was significantly smaller in the pregnant group (**p < 0.01, two-sided Mann-Whitney *U*-test). *vGluT2-Cre*, n = 19 cells from five NP females and 8 cells from five GD 15 females; *vGAT-Cre*, n = 19 cells from five NP females and 18 cells from five GD 15 females. **e**, **h**, Schematic of the virus injections. **f**, **i**, Representative coronal sections of PVH containing ChR2-YFP (**f**) and SypGFP (**i**). Blue, DAPI. Right panels show a magnified view of the white boxed areas. Scale bars, 30 μm for low magnification images and 10 μm for the magnified views. **g**, **j**, Density of axons (YFP+ pixels per μm^2^, **g**) and presynaptic puncta (# of dots per μm^2^ ×10^−3^, **j**) in PVH and DMH. *p < 0.05 and **p < 0.01 by two-sided Mann-Whitney *U*-test. N = 5–6 mice each. Error bars, SEM. See Extened Data Figs. 1–3 for more data.

Consistent with these observations, when ChR2-targeted MPN neurons were activated without restricting cell types, the net amplitude of light-evoked responses in PVH OT neurons was significantly smaller in GD 15 females (Extended Data Fig. 2m–o). Thus, the overall impact of circuit changes was biased toward a reduced excitatory drive toward the PVH OT neurons. Conversely, neither the connection ratio nor the response amplitude from *vGluT2*+ or *vGAT*+ neurons in the LHA to PVH OT neurons differed between NP and GD 15 females (Extended Data Figs. 1i, j, and 2), except for a slight reduction in the connection ratio of excitatory neurons. These observations largely align with the rabies virus tracing data, reinforcing the regional specificity of the reduced input to PVH OT neurons. Moreover, they demonstrated that this reduction was driven specifically by diminished excitatory rather than inhibitory inputs from the MPN in pregnant females. Of note, consistent with the reversible nature of this connectivity change (Fig. 1f), both the connection ratio from MPN *vGluT2*+ neurons to PVH OT neurons and the amplitude of ChR2-evoked EPSCs were restored to NP levels by postpartum day (PPD) 1 (Extended Data Fig. 1k–m).

Next, we aimed to directly visualize the axons and presynaptic structures to determine whether the reduction in excitatory MPN output was specific to the PVH or extended more broadly to other brain regions. We virally targeted either ChR2-EYFP or synaptophysin conjugated to GFP (SypGFP)^19^ in MPN *vGluT2+* neurons. Both axonal projections and presynaptic puncta originating from excitatory MPN neurons exhibited a significant reduction in density within the PVH of GD 15 females compared to NP females (Fig. 2e–i). Quantitative analysis revealed that this decrease was not evident in most other MPN output regions, such as the DMH (Fig. 2g, j, and Extended Data Fig. 3), with one notable exception: the ventromedial hypothalamus (VMH), which also showed reduced puncta density in GD 15 females (Extended Data Fig. 3i). Overall, the reduction in presynaptic puncta was more substantial than the decline in axonal labeling, suggesting that synaptic remodeling predominantly occurs at the level of local synaptic structures.

In summary, these results indicate that the female brain undergoes transient synaptic remodeling within specific pre- and post-synaptic pairs during pregnancy, which contributes to a reduced excitatory drive to PVH OT neurons.

## Feeding-related activity of PVH OT neurons declines during pregnancy

What is the biological relevance of neural circuit remodeling? Initially, we focused on the maternal functions of OT, such as parturition and lactation^20^. To counteract the reduced excitation of PVH OT neurons, we chemogenetically activated these neurons for 3 consecutive days during late pregnancy; however, this intervention did not clearly impact pregnancy duration, parturition outcomes as assessed by the number of pups born, or postnatal pup growth (Extended Data Fig. 4a–e). Accumulating evidence suggests that PVH OT neurons are more important in appetite regulation than was previously recognized. Notably, conditional knockout (cKO) of the *OT* gene in the PVH or selective ablation of PVH OT neurons in male mice induces hyperphagic obesity^21,22^. We further confirmed that *OT* cKO in female NP mice resulted in a significant increase in daily food intake and body weight (Extended Data Fig. 4f–i). Because pregnant mammals naturally increase their food intake to support fetal growth and prepare for the heightened energy demands of lactation^23–25^, we hypothesized that reduced excitation of the PVH OT neurons facilitates enhanced food intake during pregnancy.

A key prediction of our hypothesis is that feeding-related activity in PVH OT neurons diminishes during pregnancy. To evaluate this, we conducted chronic, single-cell-resolution microendoscopic Ca^2+^ imaging using GCaMP6s^26^ to monitor individual PVH OT neurons in freely moving NP and GD 16 female mice (Fig. 3a, b, Extended Data Fig. 5). Each imaging field included ∼20 regions of interest (ROIs) corresponding to single PVH OT neurons (Fig. 3c). After food presentation and feeding onset, these ROIs displayed diverse response patterns (Fig. 3d, Movie S1). Using k-means clustering based on the area under the receiver-operating-characteristic curve (auROC; Extended Data Fig. 5e, f), we grouped the responses into four responsive clusters (Clusters 1–4) and one non-responsive cluster (Cluster 5) (Fig. 3e). The responsive clusters were designated constantly responding (Cluster 1), transiently responding and decaying (Cluster 2), gradually suppressed (Cluster 3), and constantly suppressed (Cluster 4). In NP mice, roughly 50 % and 30 % of ROIs were elevated and suppressed, respectively, during feeding, indicating robust engagement of PVH OT neurons. In GD 16 mice, the fractions with elevated responses (Clusters 1 and 2) tended to decrease, whereas those with suppressed responses (Clusters 3 and 4) increased (Fig. 3f), yielding a significant reduction in overall population activity (Fig. 3g, h). This attenuation was further corroborated in ROIs reliably tracked across NP and GD 16 stages (Fig. 3i, j). Collectively, these data demonstrate marked suppression of feeding-related PVH OT neuron activity during late pregnancy.

**Fig. 3.**
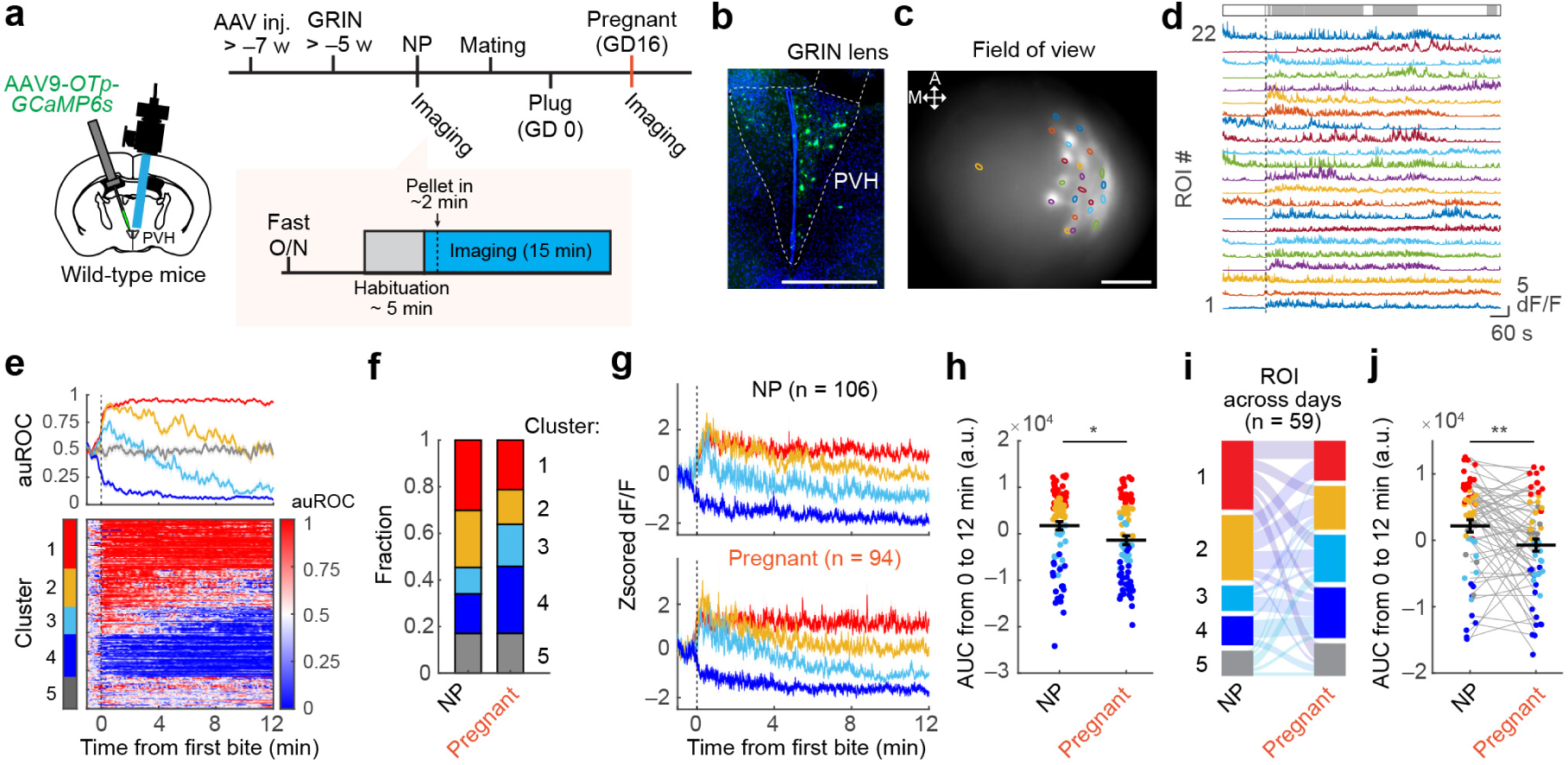
Dynamics of feeding-related activity in PVH OT neurons during pregnancy. **a**, Schematic of the experimental timeline. **b**, Representative image showing the GRIN lens tract and GCaMP6s expression (green) in the PVH. Nuclei are counterstained with DAPI (blue). Scale bar, 500 μm. **c**, Example spatial map showing ROIs overlaid on a maximum intensity projection image from a NP mouse. A, anterior; M, medial. Scale bar, 100 μm. **d**, Representative Ca^2+^ responses from individual ROIs during feeding behavior. Colors correspond to those in panel **c**. The dotted line indicates the time of the first bite. The top raster plot (gray) indicates feeding behavior. **e**, Classification of ROIs into four and one responsive and non-responsive clusters, respectively, based on response selectivity (auROC; see Methods). Top: population-averaged auROC traces for each cluster. Bottom: heat maps of auROC values for individual ROIs. n = 200 ROIs from 5 mice. **f**, Proportions of ROIs assigned to each cluster. **g**, Average Ca^2+^ response traces during feeding in NP (top) and Pregnant (bottom) states. Colors indicate cluster identities. **h**, Quantification of the area under the curve (AUC) of z-scored dF/F from 0 to 12 min. *p < 0.05 by Welch’s *t*-test. **i**, Sankey diagram showing retention or transition of cluster identities in 59 cells reliably tracked across NP and pregnant stages. **j**, Quantification of the AUC of z-scored dF/F from 0 and 12 min. **p < 0.01 by two-sided paired *t*-test. See Extended Data Figs. 5, 6 for more data.

## MPN-to-PVH circuit can suppress increased appetite during pregnancy

Finally, we examined the potential relationship between the reduced excitation of PVH OT neurons during pregnancy and feeding regulation. To this end, we first chemogenetically activated the PVH OT neurons and measured their subsequent food intake (Extended Data Fig. S6a, b). This manipulation significantly reduced food intake within the 24-h period following CNO injection, with late-pregnant females exhibiting a more pronounced reduction (Extended Data Fig. 6c, d). To evaluate the longitudinal effects of PVH OT neuron activation on feeding and body weight, we applied the same chemogenetic activation for three consecutive days during GD 14–16 or the corresponding period in NP controls (Fig. 4a, b). This treatment did not significantly alter food intake or body weight in NP females (Fig. 4c–e), which was consistent with the weaker anorexigenic effect observed in the short-term assay (Extended Data Fig. 6d). Conversely, the same procedure in pregnant females significantly reduced food intake to levels comparable to those observed in NP controls, resulting in a significant reduction in maternal body weight (Fig. 4c–e). Thus, activation of PVH OT neurons selectively counteracts pregnancy-associated enhanced feeding.

**Fig. 4.**
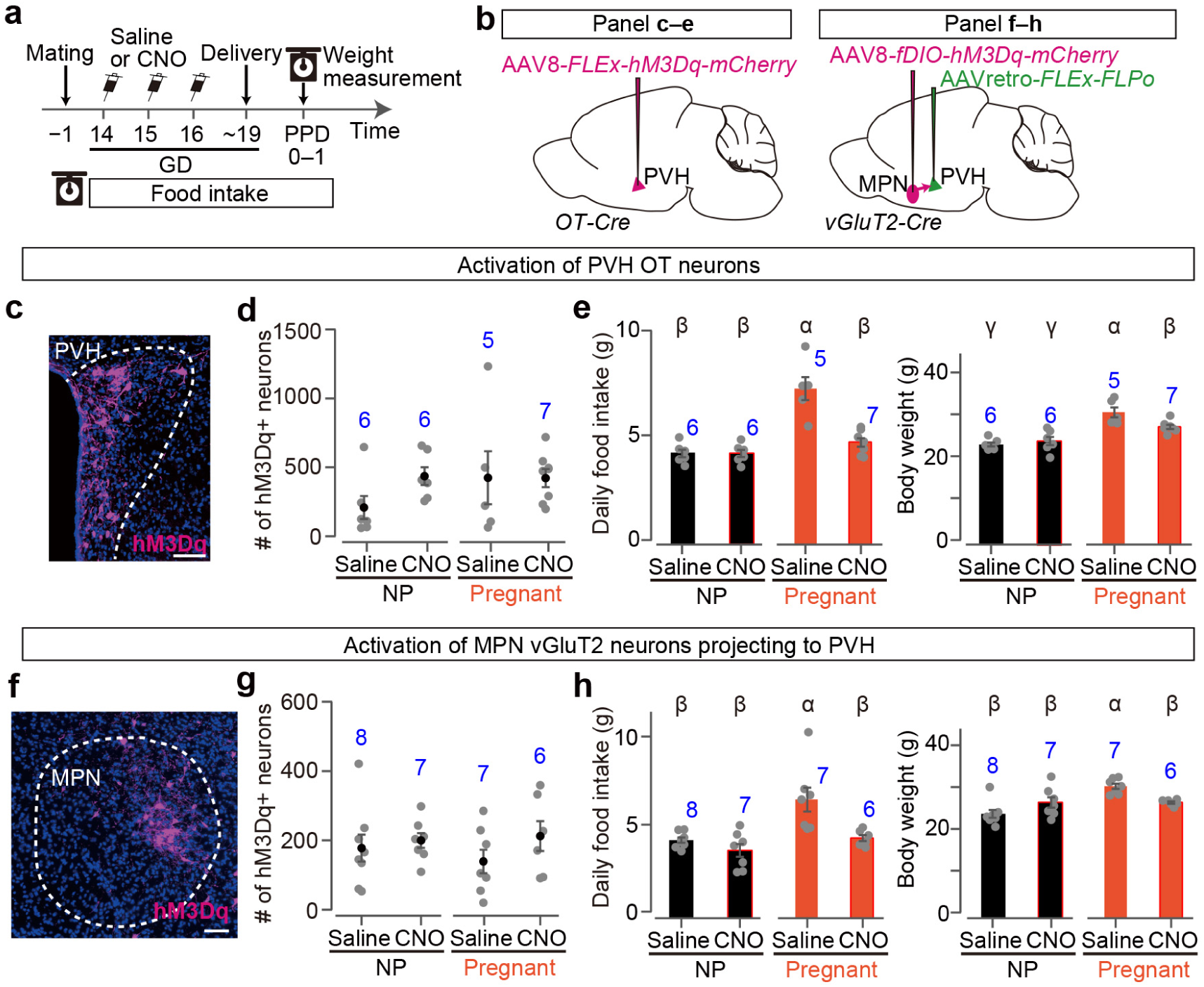
MPN-to-PVH circuit can suppress increased appetite during pregnancy. **a**, Timeline of the experiment. Saline or CNO was injected over 3 consecutive days from GD 14 to 16 in both the pregnant group and the corresponding days in the NP control group. **b**, Schematic of the virus injections. **c**, **f**, Representative coronal sections of the PVH (**c**) or MPN (**f**) expressing hM3Dq (magenta). Blue, DAPI. Scale bars, 50 μm (**c**), and 30 μm (**f**). **d**, **g**, The number of neurons expressing hM3Dq in the PVH (**d**) or MPN (**g**). No significant difference was found by one-way ANOVA. **e**, **h**, Chemogenetic activation of PVH OT neurons (**e**) or PVH-projecting MPN vGluT2+ neurons (**h**) suppressed food intake and body weight in the pregnancy phase. Different letters (α–γ) in the upper part of the graph denote significant differences at p < 0.05 by one-way ANOVA with post-hoc Tukey HSD. Error bars, SEM. The number of animals (N) is indicated in blue. See Extended data Fig. 6 for more data.

To examine the contribution of PVH-projecting MPN *vGluT2*+ neurons to feeding and body weight regulation, we performed pathway-specific chemogenetic activation during GD 14–16 (Fig. 4b), which reduced food intake and body weight only in pregnant females (Fig. 4f–h). Moreover, this activation significantly increased the PVH OT neuron activity while simultaneously reducing food intake (Extended Data Fig. 6e–h), confirming that MPN excitatory inputs can drive PVH OT neuron activity to suppress feeding. In contrast, chemogenetic activation of the PVH-projecting MPN *vGAT*+ inhibitory neurons did not affect food intake or body weight in either the NP or pregnant females (Extended Data Fig. 6i–l). Thus, these results indicate that activation of PVH-projecting MPN excitatory neurons selectively suppresses pregnancy-associated enhanced feeding. Combined with the observed pregnancy-specific decline in PVH OT neuron activity during feeding (Fig. 3), our data support a model in which reduced excitatory connectivity from the MPN to PVH OT neurons contributes to the enhanced appetite characteristics of pregnant females (see Extended Data Fig. 7 for graphical abstract).

## Discussion

What advantages might circuit-level remodeling confer during pregnancy? We observed that MPN excitatory neurons undergo targeted synaptic elimination in the PVH while preserving most other outputs (Extended Data Fig. 3), demonstrating pathway selectivity. Likewise, PVH OT neurons preferentially reduce excitatory input from the MPN yet retain inputs from many other regions, such as the LHA (Fig. 2), revealing input-origin specificity. Such dual specificities allow postsynaptic neurons to fine-tune particular functional pathways while preserving essential others. OT itself exerts functions beyond appetite suppression, notably promoting parental behaviour, social memory formation, and stress regulation^9,27^, which may manifest during pregnancy.

Decades of studies in rodents have highlighted the significance of the interaction between the arcuate nucleus (ARH) and PVH in regulating food intake^28–31^. Within the PVH, multiple neuronal populations contribute to appetite suppression, including those expressing glucagon-like peptide-1 and melanocortin-4 receptors^30,32–34^. Additionally, OT neurons are potent anorexigenic regulators^21,22,35^ and may be suppressed by hunger-encoding ARH *Agrp*+ neurons^36^. Nonetheless, a previous study failed to observe food intake–related activity in PVH OT neurons using fiber photometry^34^, raising questions regarding their physiological relevance. Our findings may explain this discrepancy. The PVH OT neuron responses during feeding were highly heterogeneous; specifically, while some neurons were activated, a substantial subset exhibited suppressed activity (Fig. 3). This bidirectional pattern may lead to the cancellation of net signals when measured using bulk calcium imaging, underscoring the importance of single-cell resolution analyses. Moreover, the conditional knockout data (Extended Data Fig. 4) and pharmacogenetic manipulations (Fig. 4) further reinforced the potent anorexigenic role of PVH OT signaling in females.

The MPN-to-PVH pathway has received relatively less attention compared to the well-characterized ARH-to-PVH pathway for feeding regulation^29–31^, highlighting the efficacy of our approach in identifying relevant presynaptic neurons without prior assumptions. The preoptic area, including the MPN, has been intensively studied in various contexts^37^, including thermal homeostasis^38,39^, parental behaviors^40^, and stress responses^41^. Recent studies have suggested that medial preoptic area neurons form monosynaptic connections with ARH *Agrp+* neurons^42^, and that certain excitatory subtypes exhibit feeding-related responses in addition to thermal inputs^43^, influencing feeding regulation through downstream PVH neurons^43^. Further research to elucidate how appetite during pregnancy is regulated through the complex interplay among the preoptic area, ARH, and PVH at the level of specific cell types is warranted.

Considering OT’s established roles in stimulating labor and milk ejection^44^, how can its dual actions—support of maternal functions and suppression of appetite—coexist within PVH OT neurons? We propose three non-mutually exclusive explanations. First, the temporal demands differ: excitatory input to PVH OT neurons is reduced specifically in late pregnancy, then restored during the periparturition period (Fig. 1f, Extended Data Fig. 1k–m), when pulsatile activity necessary for parturition and lactation emerges^45^. This suggests that distinct mechanisms enhance appetite during late pregnancy and lactation. During late pregnancy, females need to maintain a high appetite despite extremely high plasma leptin levels^23,24,46^, which may require a specialized mechanism beyond the documented leptin resistance^47–49^, such as that identified in this study. Conversely, the heightened appetite observed during lactation may be linked to reduced plasma leptin levels and increased energy demands accompanied by elevated *Agrp*+ neuron activity in ARH^50^. Second, the distinct subtypes of PVH OT neurons may play different roles. For instance, labor induction and milk ejection are facilitated by magnocellular PVH OT neurons projecting to the pituitary gland^44^, whereas the anorexigenic effects of OT may involve parvocellular PVH OT neurons projecting to the hindbrain and pons^39,51^. Future investigations employing precise genetic manipulation of PVH OT subtypes are required to delineate their functional heterogeneity. Distinct OT secretion patterns may recruit discrete downstream effectors that drive myoepithelial contractions in the uterus and mammary glands, and modulate appetite. Myoepithelial contraction depends on pulsatile, phasic bursts of OT neuron activity^44,45^, whereas feeding-associated responses are predominantly tonic (Fig. 3). Accordingly, forthcoming studies should test whether downstream effectors can discriminate between these temporal signatures of OT release.

Although the present study focused on PVH OT neurons, pregnancy-associated neural circuit remodeling is likely to extend beyond this pathway, as exemplified by the MPN-to-VMH circuit (Extended Data Fig. 3i). Because both the PVH and VMH contain diverse cell types with distinct functions, whether specific neural subpopulations within these brain regions are affected during pregnancy should be determined. Prior reports have reported reduced secretion of arginine vasopressin and corticotropin-releasing hormones from the PVH in late-pregnant rodents^23^. Therefore, we anticipate that the present approach will have broad applicability for characterizing neural circuit-level plasticity beyond the context of feeding regulation. Regarding the underlying molecular mechanisms, recent research highlighted the prominent role of sex-steroid hormone signaling in driving structural and functional plasticity in the preoptic and hypothalamic regions of female mice. For instance, estrogen receptor 1 has been shown to regulate the estrus cycle-dependent periodic remodeling of VMH-to-AVPV neural connections^11^. Similarly, MPN galanin+ neurons, the hub for parental behavior, undergo morphological and functional alterations during pregnancy in an Esr1- and progesterone receptor-dependent manner^2^. Thus, elucidating the precise molecular mechanisms governing pregnancy-associated circuit remodeling will not only deepen our understanding of adult neuroplasticity but may also offer potential strategies for manipulating neural circuits to address various brain disorders and aging.

## Materials and Methods

### Animals

The animals were housed under a 12-h light/12-h dark cycle with *ad libitum* access to food and water. Wild-type C57BL/6J mice were purchased from Japan SLC. *vGluT2-Cre* (also known as *Slc17a6-ires-Cre*, Jax #028863), *vGAT-Cre* (also known as *Slc32a1-ires-Cre*, Jax #028862), and *OT-Cre* (Jax #024234) were obtained from Jackson Laboratory. *OT* cKO in the PVH was achieved by crossing *OT^flox/flox^*and *OT^−/−^* mice, as previously described^14^. We chose the *OT^flox/–^* model to increase the cKO efficiency. Notably, if we had used *OT^flox/flox^*mice for cKO, a small fraction of the *flox* alleles that did not experience recombination would have easily masked the phenotypes due to the high expression of the *OT* gene. We used the F1 hybrid of C57BL/6 and FVB strain for microendoscopic imaging experiments^26^ (Fig. 3, Extended Data Fig. 5, and Extended Data Fig. 6 e–h). Wild-type FVB mice were purchased from CLEA Japan, Inc. (Tokyo, Japan). All experimental procedures were approved by the Institutional Animal Care and Use Committee of the RIKEN Kobe Branch.

### Viral preparations

We obtained the following AAV vectors from Addgene (titer is shown as genome particles [gp] per ml): AAV serotype 9 *hSyn-Cre* (#105555, 2.3 × 10^13^ gp/ml), AAV serotype 8 *hSyn-FLEx-hM3Dq-mCherry* (#44361, 2.1 × 10^13^ gp/ml), and AAV serotype 5 *Ef1a-FLEx-hChR2(H134R)-EYFP* (#20298, 2.5 × 10^13^ gp/ml). AAV serotype 8 *hSyn-fDIO-hM3Dq-mCherry* (#1457-aav8, 2.0 × 10^13^ gp/ml) and AAV serotype 9 *Ef1a-fDIO-ChR2(H134R)-EYFP* (#1391-aav9, 2.0 × 10^13^ gp/ml) were purchased from the Canadian Neurophotonics Platform viral vector core facility. AAV serotype 9 *OTp-mCherry* (2.9 × 10^13^ gp/ml) has been described previously^14^. The AAV vectors were generated by the University of North Carolina vector core using the plasmids: AAV serotype 5 *CAG-FLEx-TVA-mCherry* (2.4 × 10^13^ gp/ml)^15^ and AAV serotype 8 *CAG-FLEx-RG* (1.0 × 10^12^ gp/ml)^15^. *pAAV*-*CAG-FLEx-SypGFP* was constructed by PCR-based subcloning of *SypGFP* cassette^19^ into the *pAAV-CAG-FLEx* backbone (AscI/SalI fragment of *pAAV-CAG-FLEx-RG*; Addgene #48333). AAV serotype 2 *CAG-FLEx-SypGFP* (1.2 × 10^12^ gp/ml) was generated by the UNC viral core and was a gift from Koki Emori and Kazushige Touhara. *CAG-FLEx-FLPo* was derived by in-fusion-based PCR cloning using the following two DNA fragments: i) SalI-AscI restriction fragment of pAAV *CAG-FLEx-TCb* (#48332; Addgene) as a vector backbone, and ii) PCR fragment of *FLPo* originating from *pTCAV-FLEx-FLPo* (#67829; Addgene)^52^. Because a nuclear localization signal (NLS) was not included in this version of the *FLPo* cassette, we subsequently performed PCR-based subcloning of the *SV40 NLS* to the 5’ end of the *FLPo* cassette, resulting in the construction of *pAAV CAG-FLEx-FLPo*. With this plasmid, the AAV retrograde serotype *CAG-FLEx-FLPo* (3.8 × 10^12^ gp/ml) was generated by the viral vector core of the National Institute for Physiological Sciences.

### Stereotactic injection

To target the AAV in a specific brain region, stereotactic coordinates were defined for each brain region using the Allen Mouse Brain Atlas^53^. The mice were anesthetized with 65 mg/kg ketamine (Daiichi Sankyo) and 13 mg/kg xylazine (X1251, Sigma-Aldrich) via intraperitoneal injection, and then head-fixed to a stereotactic apparatus (Narishige). The following coordinates were used (in mm from the bregma for anteroposterior [AP], mediolateral [ML], and dorsoventral [DV]): PVH, AP −0.8, ML 0.2, DV 4.5; LHA, AP −1.7, ML 1.0, DV 5.2; MPN, AP −0.2, ML 0.2, DV 5.2. The volume injected to express ChR2 (AAV*-FLEx-ChR2* or a mixture of AAV*-FLEx-ChR2* and AAV*-Cre*) was 100 nl at a speed of 50 nl/min. The injection volume and speed of the other viruses were 200 nl and 50 nl/min, respectively. Following viral injection, the animals were returned to their home cages.

### Trans-synaptic retrograde tracing

The rabies virus was prepared using the following protocol^54^ with viruses, cell lines, and materials, as previously described^14^. Briefly, RV*d*G-GFP was prepared using the B7GG cell line (a gift from Ed Callaway) and plasmids (*pCAG-B19N*, *pCAG-B19P*, *pCAG-B19L*, *pCAG-B19G*, and *pSADdG-GFP-F2*; gifts from Ed Callaway), and then pseudotyped using BHK-EnvA cells (a gift from Ed Callaway). The titer of RV*d*G-GFP+EnvA used in this study was estimated to be 4 × 10^9^ infectious particles per ml based on serial dilutions of the virus stock, followed by infection of the HEK293-TVA800 cell line (a gift from Ed Callaway).

For trans-synaptic tracing using the rabies virus, 200 nl of a 1:1 mixture of AAV5 *CAG-FLEx-TVA-mCherry* and AAV8 *CAG-FLEx-RG* was injected into the unilateral PVH of *OT-Cre* mice. A virgin female (7–13 weeks old) that had spent 2 weeks after the injection of AAV*-FLEx-TVA-mCherry* and AAV*-FLEx-RG* was mated with a wild-type male. The following day, the male was removed from the cage and the vaginal plug was checked. GD 0 was defined as the day on which the vaginal plug was detected. The females that failed to form a plug were classified as “non-pregnant (NP).” In Fig. 1a–d, 200 nl of RV*dG-GFP*+EnvA was injected into the same brain region to initiate trans*-*synaptic tracing on GD 13. One week after injection of the rabies virus, the mice were sacrificed and perfused with phosphate-buffered saline (PBS), followed by 4% paraformaldehyde (PFA) in PBS. In Fig. 1e, f, the rabies virus was injected on the day indicated in Fig. 1e, and the mice were sacrificed 1 week after the injection.

Quantitative analysis of trans-synaptic tracing samples was performed by experimenters who were blinded to the results of the electrophysiology and food intake assays. Because a negative control experiment omitting Cre expression showed nonspecific rabies labeling^15^ in the PVH and paraventricular thalamus^45^, we excluded these regions from the analysis. Fluorescent protein-expressing neurons were counted as previously described^45^. Briefly, 30 μm coronal sections were collected, and every third section was subjected to cell counting. Each section was imaged using a slide scanner (Axio Scan.Z1, Zeiss) with a 20× objective lens (N.A. 0.8). Image processing was performed semi-automatically using the ImageJ macro (National Institutes of Health). Briefly, annotators blinded to the experimental conditions manually delineated regions of interest (ROIs) for each brain area on the DAPI channel—which visualizes brain structures but not GFP⁺ cells— using the Allen Brain Atlas^53^. For each ROI, raw fluorescence images of GFP or mCherry were processed with an open filter and a median filter. Labeled cells were then identified with the “Threshold” and “Analyze Particles” commands. Starter cells (Fig. 1b) were defined as those detected in both GFP and mCherry channels. Because every third section was analyzed, the number of starter cells reported in Fig. 1b was adjusted by a factor of 3. In Fig. 1d, the d′ value was calculated as

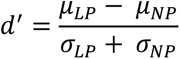

where μ and σ denote the average and standard deviations, respectively. In Fig. 1f, the d′ value was calculated as above, but “LP” should be replaced with each GD or PPD.

### Electrophysiology

Electrophysiological recordings were performed by an experimenter blinded to the results of the rabies virus-mediated trans-synaptic tracing and food intake assays. Females were used for electrophysiological experiments after 4 weeks of ChR2 expression following AAV injection and an additional 2 weeks for mCherry expression. The targeting specificity of OTp-mCherry has been established previously^14^. At GD 15, electrophysiological recordings were performed as described previously^14^. Briefly, mice were deeply anesthetized with isoflurane. Acute coronal slices (250-μm thick) were prepared using a linear slicer (Pro7N, Dosaka EM) in an ice-cold slicing solution containing (in mM): 230 sucrose, 3 KCl, 1.2 KH_2_PO_4_, 10 MgSO_4_, 10 D-glucose, 26 NaHCO_3_, 2 CaCl_2_, 5 Na ascorbate, and 3 Na pyruvate. The slices were incubated and maintained at room temperature for at least 30 min before recording in a solution containing (in mM): 92 NaCl, 20 HEPES, 2.5 KCl, 30 NaHCO_3_, 1.2 NaH_2_PO_4_, 2 CaCl_2_, 2 MgSO_4_, 25 D-glucose, 5 Na ascorbate, and 3 Na pyruvate (pH ∼7.3; osmolarity adjusted to 295–305 mOsm). During the recording, individual slices were perfused with ACSF containing (in mM): 126 NaCl, 26 NaHCO_3_, 2.5 KCl, 1.25 NaH_2_PO_4_, 1 MgSO_4_, 12.5 D-glucose, and 2 CaCl_2_ (pH ∼7.3, osmolarity adjusted to 295–305 mOsm). NBQX (0373, Tocris) or gabazine (SR95531, Tocris) were dissolved at 10 μM in the ACSF. All solutions were continuously bubbled with 95% O_2_/5% CO_2_. The bath temperature was monitored and adjusted to 32–34 °C (TC-324B; Warner Instruments). Patch pipettes having a resistance of 4–6 MΩ were fabricated from thin-wall glass capillaries (1.5 mm o.d./1.12 mm i.d., TW150F-3, World Precision Instruments). The internal patch pipette solution used for voltage-clamp recordings contained (in mM): 117 cesium methanesulfonate, 20 HEPES, 0.4 EGTA, 2.8 NaCl, 0.3 Na_3_GTP, 4 MgATP, 10 QX314, 0.1 spermine, and 13 biocytin hydrazide (pH ∼7.3, osmolarity adjusted to 285–295 mOsm). For the current-clamp experiments, the patch pipettes were filled with a solution containing (in mM): 126 K gluconate, 10 HEPES, 0.3 EGTA, 4 KCl, 10 phosphocreatine disodium salt hydrate, 0.3 Na_3_GTP, 4 MgATP, and 13 biocytin hydrazide (pH ∼7.3, osmolarity adjusted to 285–295 mOsm). Electrophysiological recordings were performed using a Multiclamp 700B amplifier (Molecular Devices), with a low-pass filter at 1 kHz and digitization at 10 kHz. In voltage-clamp mode, cells were held at −70 mV or 0 mV. In current-clamp mode, cells were held at around −70 mV by injecting a hyperpolarizing current.

Optogenetic stimulation (1 ms) was achieved using a 470-nm LED light (M470L3, Thorlabs) controlled by a pulse generator (Master-8, AMPI). The optical intensity of the LED light was adjusted to 3.6 mW measured at the back aperture of an objective lens (S120VC sensor; Thorlabs). PVH OT neurons were targeted using mCherry fluorescence. The connected ratio of an animal (Fig. 2c, d) was calculated as follows: from each animal, we recorded from five or more OT neurons and the recorded OT neuron was regarded as “connected” to the stimulated neurons if the cell showed a time-locked response within 10 ms after a light pulse in eight or higher trials out of 10 trials. The response amplitude was obtained by subtracting the baseline (an average of 50 ms before the light pulse) from the average of 5 ms around the peak. The onset of the response and sEPSCs/sIPSCs were detected using a previously described algorithm^55^. In Extended Data Fig. 1, physiological properties were measured shortly after attaining the whole-cell configuration. In Fig. 2c, d, and Extended Data Fig. 1l, i, j, and 2n, each trace is an average of 10 trials.

### Histochemistry

The mice were anesthetized with isoflurane and perfused with PBS, followed by 4% PFA in PBS. The brains were post-fixed overnight with 4% PFA. Twenty-μm coronal brain sections were made using a cryostat (Leica). Fluorescence *in situ* hybridization was performed as previously described^14,56^. The primers (5’ – 3’) to produce RNA probes were: *OT*, forward, AAGGTCGGTCTGGGCCGGAGA, reverse, TAAGCCAAGCAGGCAGCAAGC; *Cre*, forward, CCAAGAAGAAGAGGAAGGTGTC, reverse, ATCCCCAGAAATGCCAGATTAC. Anti-GFP (GFP-1010; Aves Labs; 1:500) and anti-RFP (5f8; Chromotek; 1:500) primary antibodies were used. Signal-positive cells were detected using anti-chicken Alexa Fluor 488 (703-545-155;Jackson ImmunoResearch; 1:500) and anti-rat Cy3 (712-165-153; Jackson ImmunoResearch; 1:500) antibodies. Fluoromount (K024; Diagnostic BioSystems) was used as the mounting medium. Brain images were acquired using an Olympus BX53 microscope equipped with a 10× (N.A. 0.4). Signal-positive cells were counted manually using the ImageJ Cell Counter plugin.

### Quantification of axons and presynaptic puncta

In Fig. 2 and Extended Data Fig. 3, AAV5-*Ef1a-FLEx-hChR2(H134R)-EYFP* or AAV2*-FLEx-SypGFP* was injected into the unilateral MPN of *vGluT2-Cre* female mice to visualize axons and presynaptic puncta, respectively. Two weeks following the injection, each mouse was crossed with a wild-type male. At GD 15, mice were sacrificed, and 20-μm coronal brain sections were made as described above. After staining with the anti-GFP antibody, brain images were acquired using an Olympus BX53 microscope equipped with a 10× (N.A. 0.4) objective lens. In Fig. 2g and Extended Data Fig. 3d, the obtained images were binarized by applying an automatic thresholding function in ImageJ. The number of white pixels (EYFP+) within the defined area was measured. In Fig. 2j and Extended Data Fig. 3i, every dot-like structure within a defined area was counted manually using the ImageJ Cell Counter plugin and reported as the number of dots per area.

### Measurement of food intake and body weight

Food intake and body weight were measured by an experimenter blinded to the working hypothesis. In Extended Data Fig. 4f–i, AAV9*-Cre* was injected into the bilateral PVH of 8-week-old *OT^flox/–^*females, and these mice were subjected to a food intake assay 3 weeks after injection. In Fig. 4c–e, AAV8*-FLEx-hM3Dq-mCherry* was injected into the bilateral PVH, and the females were subjected to a food intake assay 2 weeks or longer after the injection. In Fig. 4f–h, AAVretro*-FLEx-FLPo* was injected bilaterally into the PVH of 8-week-old female mice. Two weeks later, AAV8*-fDIO-hM3Dq-mCherry* was injected bilaterally into the MPN. Two weeks or more following the injection of the AAV that expresses *hM3Dq-mCherry*, each female mouse was used for the food intake assay. During the food intake assay, all animals were housed individually in small cages. Food intake was measured by placing pre-weighed (95 g) standard food pellets (MFG; Oriental Yeast, Shiga, Japan; 3.57 kcal/g) on the plate of the cage and reweighing them at a defined interval. Daily food intake was calculated by dividing the cumulative food intake by the corresponding number of days and reported with significance digits of 0.1 g. After measuring food intake, the body weights of the mice were measured.

### *In vivo* microendoscopic imaging

For microendoscopic recordings, AAV9-*OTp-GCaMP6s* was injected into the PVH. The targeting specificity of GCaMP6s has been established previously^26,57^. Two weeks after the viral injection, a ProView integrated GRIN lens (500 μm diameter, 8.4 mm length, cat#1050-004414, Inscopix) was inserted into the PVH. Mice were anesthetized with 65 mg/kg ketamine (Daiichi-Sankyo) and 13 mg/kg xylazine (Sigma-Aldrich) via intraperitoneal injection and head-fixed to stereotaxic equipment (RWD). A 1-mm diameter craniotomy was performed over the lens target area, centered 0.5 mm posterior to the bregma and 0.5 mm lateral to the midline. The remaining bone and overlying dura were carefully removed using fine forceps. Brain tissue was aspirated to a depth of 3 mm from the surface. The integrated GRIN lens was mounted onto the microscope and attached to the stereotaxic equipment using an Inscopix gripper. Subsequently, the lens was centered via craniotomy, and the assembly was slowly lowered into the brain, advancing the lens vertically by 4.4 mm at a 5° angle. This approach ensures that the lens center is positioned approximately 200 µm from the midline and 4.3 mm beneath the brain surface. After reaching the desired depth, the lens and baseplate were permanently fixed using SuperBond (Sun Medical) and sealed in the skull. A metal bar (CF-10; Narishige) was affixed to facilitate its easy attachment and detachment from the microscope. After the adhesive was fully attached, the microendoscope was carefully detached from the lenses. Following more than 3 weeks of recovery, the microscope was attached, and the mice were allowed to explore freely in their home cages for 5–10 min. This habituation session was conducted more than twice prior to the first imaging session.

We allowed more than 8 weeks after AAV injection and at least 6 weeks after GRIN lens implantation to ensure stable GCaMP expression. Previous studies have demonstrated that the field of view stabilizes approximately 5 weeks post-GRIN lens implantation^58^. We performed microendoscopic imaging using the Inscopix nVista system (Inscopix) without refocusing across imaging sessions within a single day, while adjusting the focal plane each day before the first imaging session. Prior to imaging, the microendoscope was attached to the animal by securing an implanted headbar. Images (1080 × 1080 pixels) were acquired using nVista HD software (Inscopix) at 10 Hz, with LED power set to 0.6–0.7 mW/mm^2^ and a gain of 3.0–5.0. The timestamps of the imaging frames and camera data were collected for alignment using WaveSurfer (https://wavesurfer.janelia.org/). Each imaging session lasted for 15 min. Following an overnight fast, neural activity was recorded for 15 min, with a food pellet introduced into the cage approximately 2 min after imaging started. The imaging data were cropped to 800 × 700 pixels and exported as .tiff files using Inscopix data processing software. To identify putative cell bodies for neural signal extraction, we used v2 of MIN1PIPE (https://github.com/JinghaoLu/MIN1PIPE^59^) with a spatial downsampling rate of 2. All traces from the identified cells were manually inspected to ensure signal quality and were excluded if they exhibited abnormal shapes or overlapped signals from adjacent cells. Relative changes in calcium fluorescence were calculated by ΔF/F_0_ = (F – F_0_)/F_0_ (where F_0_ represents the median fluorescence of the entire trace). For the longitudinal tracking shown in Fig. 3i, j, we identified the same cells across different days through manual inspection. Specifically, projection images were generated, and the detected ROIs were superimposed. By comparing these images, we matched the same cells between NP and Pregnant.

### Clustering analysis

To identify potential functional clusters and to characterize heterogeneity in feeding-related activity patterns, we conducted a clustering analysis based on the area under the receiver operating characteristic curve (auROC). For each ROI, auROC values were calculated by comparing the distribution of Ca^2+^ responses within a 100-frame window (−50 to +50 frames relative to the time point of interest; ±5 s) with the baseline distribution (600 frames recorded from −62 to −2 s relative to food-pellet entry, resampled to 100 frames to match the comparison window). The auROC ranges from 0 to 1 and quantifies the discriminability of an ideal observer: values near 0.5 indicate no selectivity, whereas values approaching 1 or 0 denote elevated or suppressed responses, respectively, during feeding.

The resulting auROC matrix (200 ROIs × 9000 frames) was subjected to principal component analysis (PCA) to reduce the dimensionality and extract components capturing the majority of the data variance. PCA was performed using the MATLAB *pca* function, yielding principal components (PCs), scores, and the associated explained variance. We then applied *k*-means clustering to the scores of the first 22 PCs, which accounted for more than 95% of the variance. Clustering was performed using the *kmeans* function in MATLAB, with five clusters and a fixed random seed for reproducibility. Each cluster displayed distinct population-averaged auROC traces and individual ROI heat maps, indicating diverse functional activity patterns in PVH OT neurons during feeding (Fig. 3e).

To quantify the AUC of the Ca^2+^ trace (Fig. 3h, j), the ROIs from the non-responsive cluster (cluster 5) were excluded. Baseline activity was defined as the period from –60 to –30 s relative to the first bite. The AUC was computed from 0 to 12 min using the MATLAB *trapz* function.

### Microendoscopic Imaging with Chemogenetic Activation of MPN Glutamatergic Neurons

For microendoscopic imaging combined with chemogenetic activation of MPN glutamatergic neurons (Extended Data Fig. 6e–h), we bilaterally injected a mixture of AAVretro-*FLEx-FlpO* and AAV9-*OTp-GCaMP6s* into the PVH, and AAV8-*FLEx-hM3Dq-mCherry* into the MPN. After 2 weeks of recovery, a GRIN lens and baseplate (Integrated Proview lens, Inscopix) were implanted above the PVH. Following an additional recovery period of at least 5 weeks, females were mated with males of the same strain, and GD 0 was defined as the day of vaginal plug detection.

On GD 15, after an overnight fast, 120 µL saline was injected intraperitoneally, and imaging began 15 min later. A food pellet was introduced 2 min after imaging began. Each imaging session lasted 15 min. After the session, the mice had ad libitum access to food for 6–7 h and were then fasted overnight. On GD 16, 120 µL clozapine-N-oxide (CNO; 0.5 mg/mL) was injected intraperitoneally, and imaging was performed exactly as on GD 15.

To quantify baseline activity before and after chemogenetic activation, Ca^2+^ traces from each ROI on GDs 15 and 16 were concatenated, and dF/F was calculated using the mean fluorescence of the entire concatenated trace (GD 15 + GD 16) as F_0_. Baseline activity was assessed during the –60 to –30 s window preceding the first bite each day (Extended Data Fig. 6g). Food intake was measured by weighing food pellets before and after the 15-min imaging session (Extended Data Fig. 6h).

### Data analysis

All mean values are reported as the mean ± SEM. The statistical details of each experiment, including the statistical tests used, the exact value of n, and the meaning of n, are provided in the figure legends. P-values are shown in each figure legend or panel; non-significant values are not noted.

## Data and materials availability

Custom MATLAB/Python scripts supporting the findings of this study and data from microendoscopy will be deposited in the SSBD depository (https://xxx). All other data are available in the main text and supplementary materials. All materials are available through requests to the corresponding authors.

## Acknowledgments

We thank Yusaku Iwasaki, Masahide Yoshida, Tatsushi Onaka, Alina Xiao, Liqun Luo, and the members of the Miyamichi laboratory for critically reading the manuscript. We also thank Koki Emori and Kazushige Touhara for providing AAV2 *CAG-FLEx-SypGFP*; Edward M. Callaway for sharing the B7GG, BHK-EnvA, and HEK293-TVA800 cell lines; Kazuko Tsujimoto for their contributions to the generation of pseudotyped rabies virus; and Kazuko Tsujimoto, Hsiao-Ling Chiang, and Satsuki Irie for their support in the data analysis.

## Funding

RIKEN Special Postdoctoral Researcher Program (KI)

RIKEN BDR-Ohtsuka Pharmaceutical Collaboration Center Kakehashi Program (KI) JSPS KAKENHI 25H02524 (KI)

JST PRESTO program JPMJPR21S7 (GT)

JSPS KAKENHI 20K15907 (HY)

JSPS KAKENHI 23H04945, 23H04939, and 25K02368 (KM)

JST CREST program JPMJCR2021 (KM)

## Author contributions

Conceptualization: KI, GT, HY, KM

Methodology: KI, GT, HY, KK

Investigation: KI, GT, HY, MH

Visualization: KI, GT, HY, KM

Funding acquisition: KI, GT, HY, KM

Supervision: KM

Writing – original draft: KI, GT, HY, KM

## Competing interests

The authors have no competing interests to disclose.

## Extended Data Figures

**Extended Data Fig. 1.**
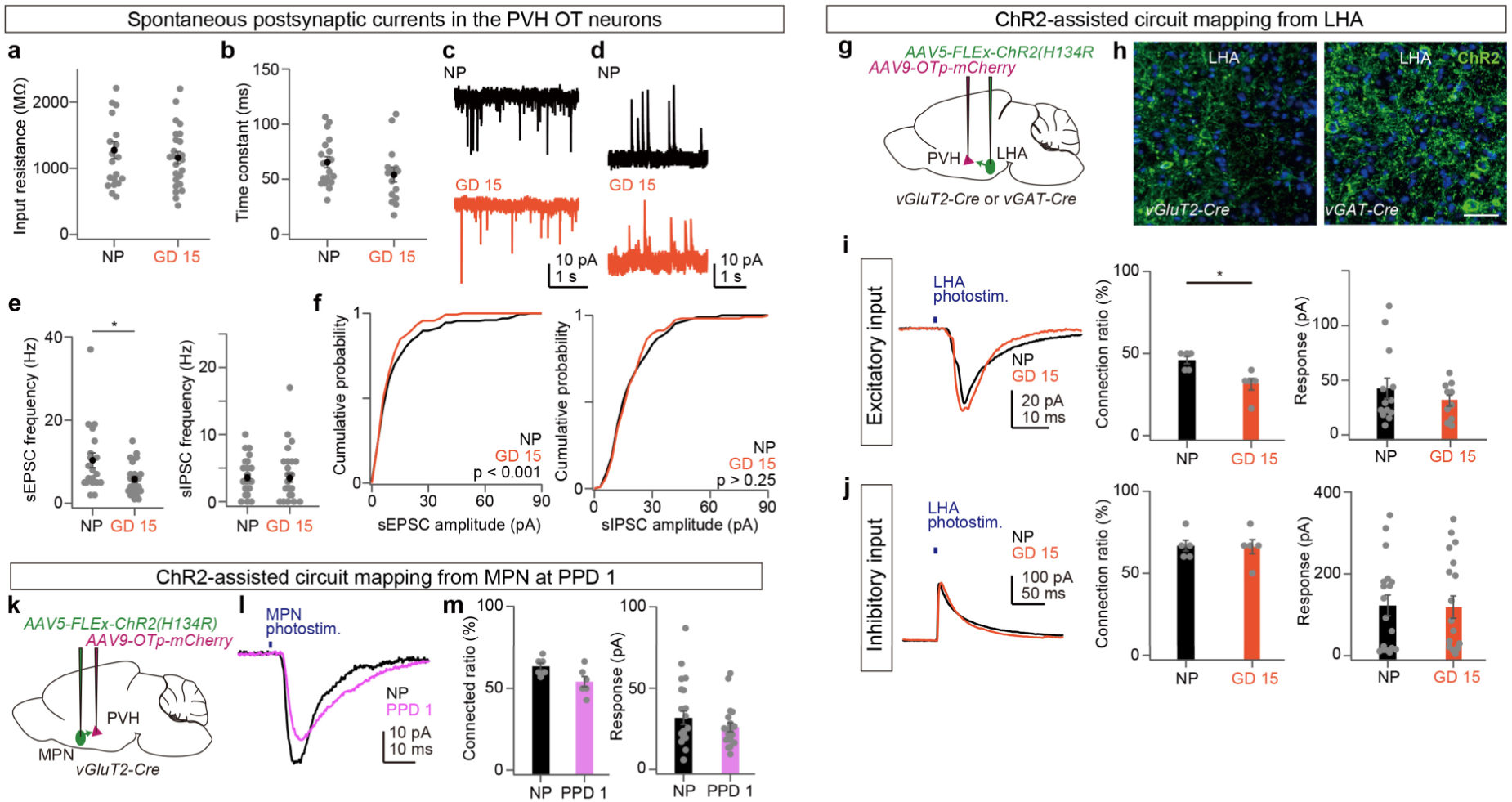
Electrophysiological properties of PVH OT neurons and additional CRACM data, related to Fig. 2. **a**, Input resistance measured at soma was not significantly different (n = 20 cells from seven NP females and 24 cells from eight GD 15 females; p > 0.80, two-tailed Student’s *t*-test). **b**, Membrane time constant was not significantly different (n = 18 cells from five NP females and 15 cells from five GD 15 females; p > 0.18, two-tailed Student’s *t*-test). **c**, **d**, Representative traces showing spontaneous excitatory postsynaptic current (sEPSC) (**c**) or spontaneous inhibitory postsynaptic current (sIPSC) (**d**) recorded from PVH OT neurons. **e**, The frequency of sEPSC is significantly lower in GD 15 females than in NP females (*p < 0.05, two-sided Student’s *t*-test), while sIPSC is not significantly different (p > 0.85). sEPSC, n = 20 cells from 7 NP females and 24 cells from 8 GD 15 females; sIPSC, n = 30 cells from 9 NP females and 29 cells from 9 GD 15 females. **f**, The distribution of sEPSC amplitude is significantly different (Kolmogorov-Smirnov test). **g**, Schematic of the virus injections for CRACM from LHA neurons. **h**, Representative coronal section of *vGluT2+* (left) or *vGAT+* (right) neurons in the LHA. Green is ChR2-EYFP amplified by anti-GFP staining. Blue, DAPI. Scale bar, 20 μm. **i**, **j**, Left: representative responses from PVH OT neurons to optogenetic activation of *vGluT2*+ (**i**) or *vGAT+* (**j**) axons originating from LHA neurons. Each trace is an average of 10 trials. Blue bar, light stimulation (1 ms, 3.6 mW). Middle: ratio showing that the recorded OT neuron was connected to the stimulated neurons. N = 5 mice each. *p < 0.05, two-sided Mann-Whitney *U*-test. Right: The light-induced response of PVH OT neurons. No significantly difference was found by the two-sided Mann-Whitney *U*-test. *vGluT2-Cre*, 13 cells from five NP females and 10 cells from five GD 15 females; *vGAT-Cre*, 18 cells from five NP females and 19 cells from five GD 15 females. **k**, Schematic of the virus injections for CRACM from MPN excitatory neurons at PPD 1 compared to the NP group. **l**, Representative responses from PVH OT neurons to optogenetic activation of *vGluT2*+ axons originating from MPN neurons. Each trace is an average of 10 trials. Blue bar, light stimulation (1 ms, 3.6 mW). **m**, Left: The ratio showing that the recorded OT neuron was connected to the stimulated neurons. N = 6 mice each. No significant difference was noted by the two-sided Mann-Whitney *U*-test. Right: The light-induced response of PVH OT neurons. No significant difference was found by the two-sided Mann-Whitney *U*-test. 23 cells from six NP females and 21 cells from six PPD 1 females. Error bars, SEM.

**Extended Data Fig. 2.**
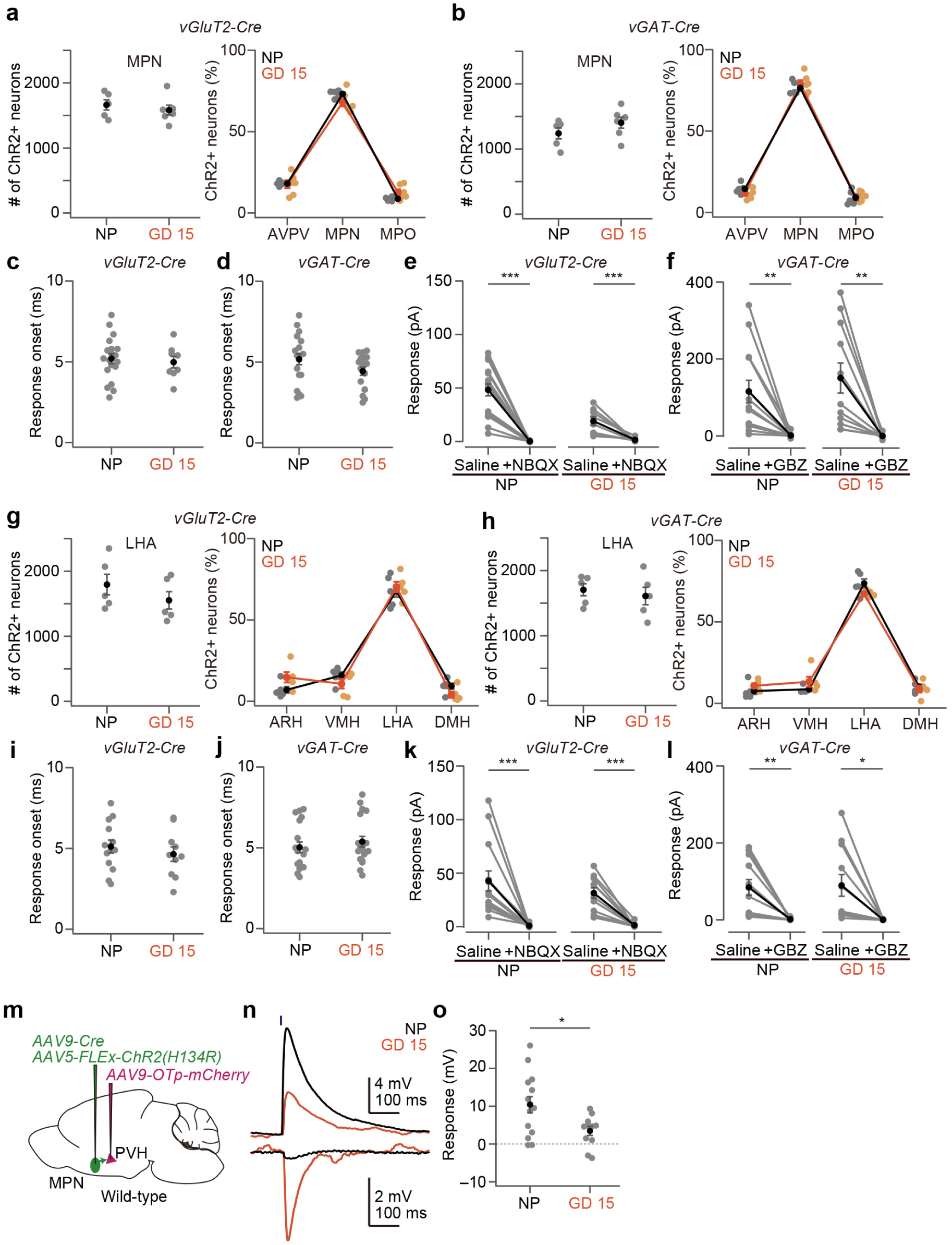
Distribution of neurons expressing ChR2 and optogenetic activation of pan-MPN neurons, related to Fig. 2. **a**, **b**, Cells expressing ChR2-EYFP were counted in the MPN (left), and a fraction of ChR2+ neurons was calculated in the MPN and two neighboring hypothalamic nuclei (right) because both *vGluT2-Cre* and *vGAT-Cre* can express ChR2 beyond the targeted area. *vGluT2-Cre*, n = 5 and 6 for NP females and LP females, respectively; *vGAT-Cre*, n = 5 mice each. **c**, **d**, Response onset of data shown in Fig. 2**c**, **d**. **e**, **f**, Optogenetic activation of *vGluT2+* and *vGAT+* MPN neurons evoked glutamatergic and GABAergic synaptic transmission, respectively (**p < 0.01, ***p < 0.001, two-tailed Welch’s *t*-test). GBZ, gabazine. *vGluT2-Cre*, n = 17 cells from five NP females and 8 cells from five GD 15 females; *vGAT-Cre*, n = 13 cells from five NP females and 11 cells from five GD 15 females. **g**, **h**, Cells expressing ChR2-EYFP were counted in the LHA (left) and a fraction of ChR2+ cells in the LHA and three neighboring hypothalamic nuclei (right). *vGluT2-Cre*, n = 5 each; *vGAT-Cre*, n = 5 mice each. **i**, **j**, Response onset of data shown in Extended Data Fig. 1i, j. **k**, **l**, Optogenetic activation of *vGluT2+* and *vGAT+* LHA neurons evoked glutamatergic and GABAergic synaptic transmission, respectively (*p < 0.05, **p < 0.01, ***p < 0.001, two-tailed Welch’s *t*-test). *vGluT2-Cre*, n = 13 cells from five NP females and 10 cells from five GD 15 females; *vGAT-Cre*, n = 13 cells from five NP females and 11 cells from five GD 15 females. **m**, Schematic of the virus injections. **n**, Representative responses of the OT neurons to light stimulation (blue bar, 1 ms, 3.6 mW). Each trace is an average of 10 trials. Note that some OT neurons exhibited inhibitory responses. **o**, Response of the OT neurons was smaller in the GD 15 females (*p < 0.05, two-sided Mann-Whitney *U*-test). n = 14 cells from seven NP females and 11 cells from GD 15 females. Error bars, SEM.

**Extended Data Fig. 3.**
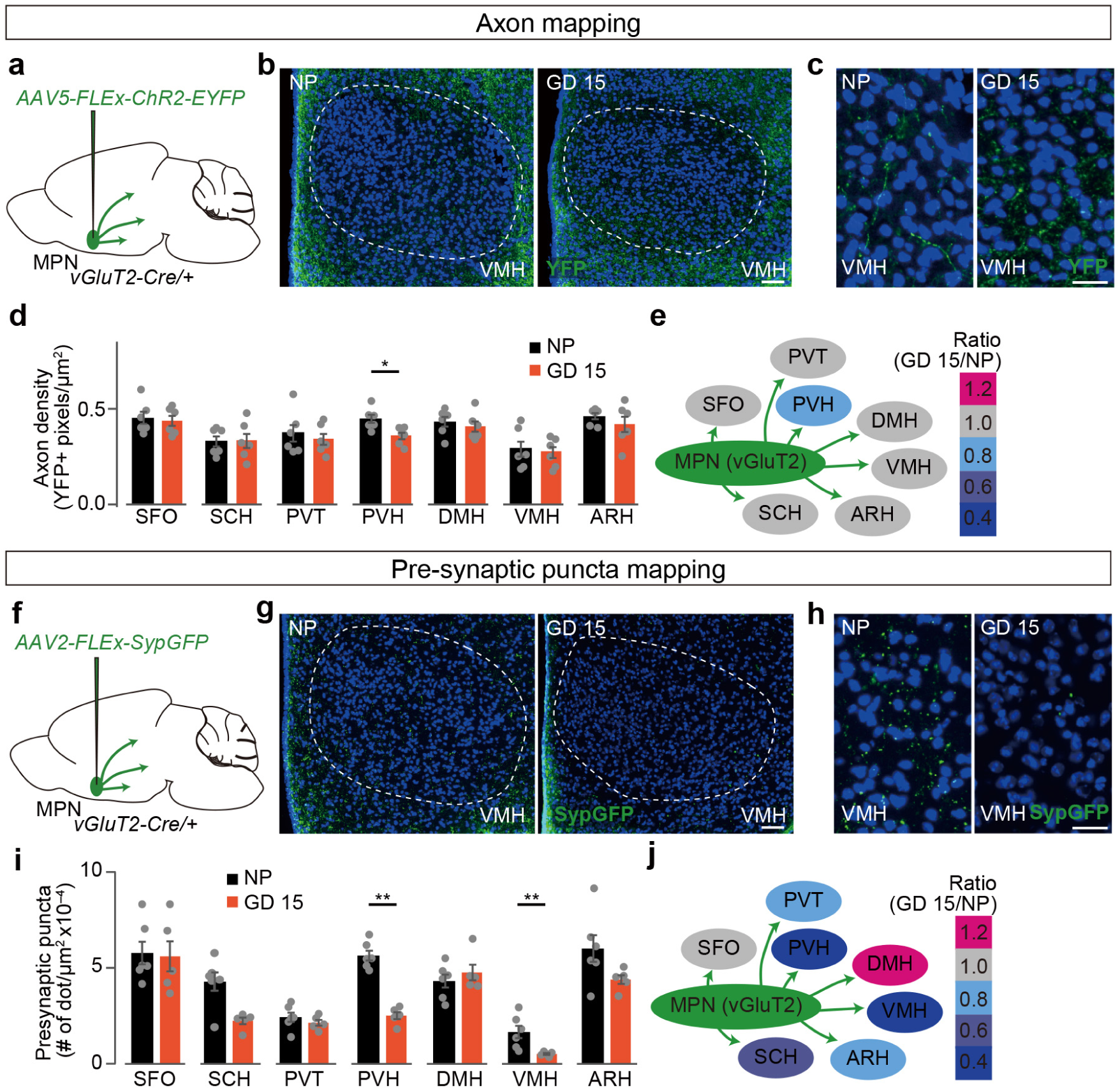
Density of axons and presynaptic puncta of *vGluT2+* MPN neurons, related to Fig. 2. **a**, **f**, Schematic of the virus injections. **b**, **g**, Representative coronal section of VMH containing ChR2-YFP (**b**) and SypGFP (**g**). Blue, DAPI. Scale bars, 30 μm. **c**, **h**, Magnified views of VMH. Scale bars, 10 μm. **d**, **i**, Density of axons (YFP+ pixels per μm^2^, **d**) and presynaptic puncta (# of dots per μm^2^ ×10^−3^, **i**) in each brain region. **p < 0.01, two-sided Mann-Whitney *U*-test. n = 5–6 mice each. **e**, **j**, Colored heat map showing fold changes of axon density (**e**) and presynaptic puncta (**j**) in each brain region. Error bars, standard error of mean (SEM).

**Extended Data Fig. 4.**
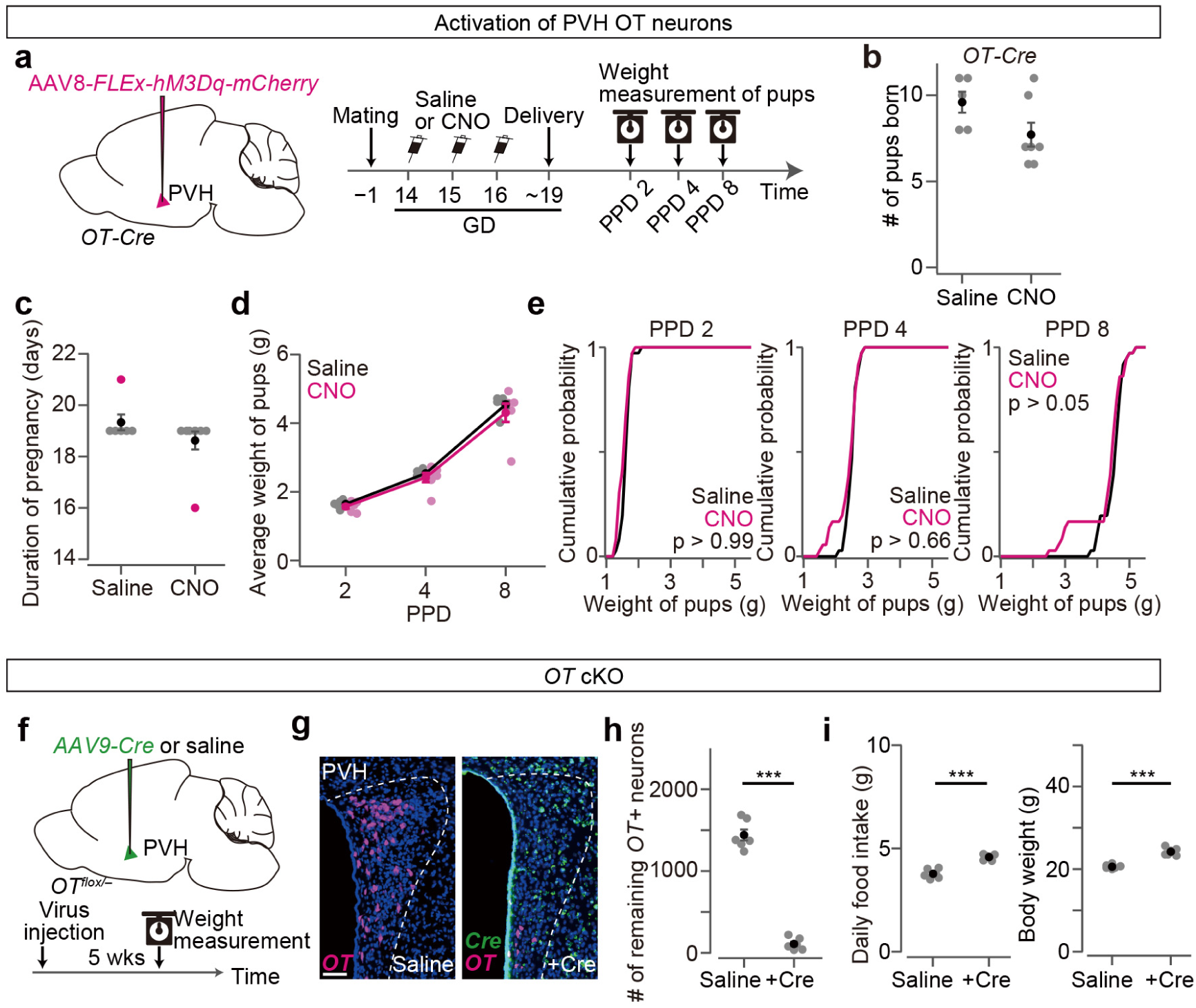
Chemogenetic activations of PVH OT neurons and *OT* cKO, related to Fig. 3. **a**, Schematic of the virus injections (left) and the timeline of the experiment (right). Saline or CNO was injected over three consecutive days from GD 14 to 16. **b**, The number of pups born. p > 0.08, two-tailed Mann-Whitney *U*-test; N = 5 and 7 for saline and CNO, respectively. **c**, The duration of pregnancy shows no statistical difference (p > 0.44, two-sided Mann–Whitney *U*-test). N = 6 and 8 mice for saline and CNO, respectively. Magenta dots indicate stillbirth. **d**, Average weight of pups per dam. No statistical difference was found in saline and CNO (two-way ANOVA with repeated measurements). N = 6 mice each. **e**, Cumulative probability of weight of pups at PPD 2, PPD 4, and PPD 8. The p-value is shown in the panel (Kolmogorov–Smirnov test). N = 6 dams each. **f**, Schematic of the virus injection (top) and the timeline of the experiment (bottom) for cKO of the *OT* gene. **g**, Representative coronal sections of the PVH from *OT^flox/−^* females without (saline; left) or with (+Cre; right) AAV*-Cre* injection. *OT* and *Cre in situ* hybridization staining are shown in magenta and green, respectively. Blue, DAPI. Scale bar, 50 μm. **h**, The number of remaining *OT*+ neurons in the PVH. N = 6 mice each. ***p < 0.001 by two-sided Welch’s *t*-test. **i**, Food intake and body weight of the pregnant and NP groups. ***p < 0.001 by two-sided Welch’s *t*-test.

**Extended Data Fig. 5.**
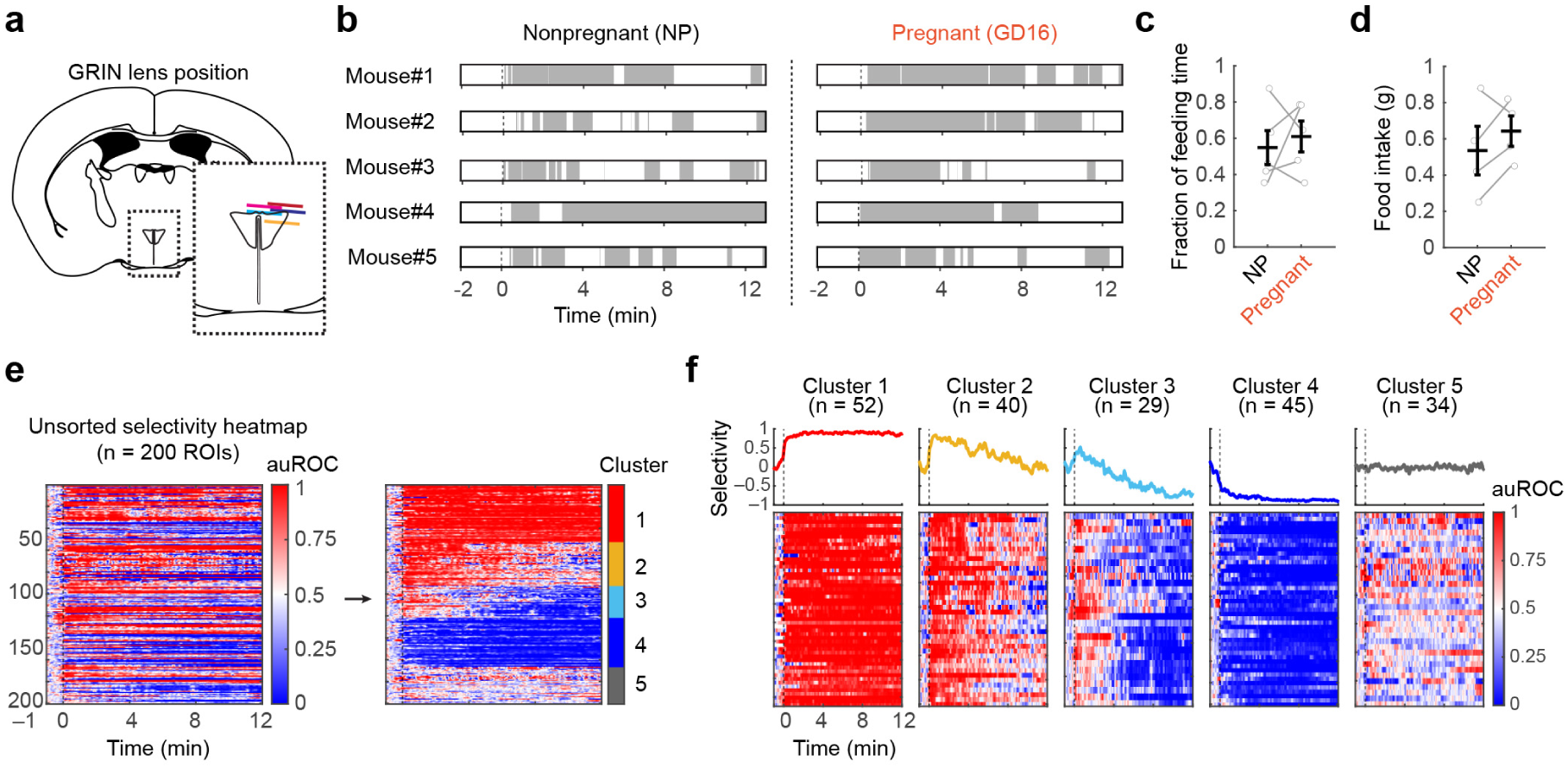
Additional analyses of microendoscopic imaging of PVH OT neurons, related to Fig. 3. **a**, Schematic of coronal sections showing the position of the GRIN lens (N = 5 mice). **b**, Raster plot (gray) indicating feeding behavior during the imaging session. Time zero corresponds to the food supply. **c**, Left: Fraction of time spent feeding. **d**, Total food intake during the imaging session. ns, not significant by two-sided Welch’s *t*-test. **e**, Heat maps showing auROC values for individual ROIs before (left) and after (right) *k*-means clustering. n = 200 ROIs from 5 mice. Time = 0 indicates the first bite. **f**, Population-averaged auROC traces and corresponding heat maps of auROC values for each cluster. Time = 0 indicates the first bite.

**Extended Data Fig. 6.**
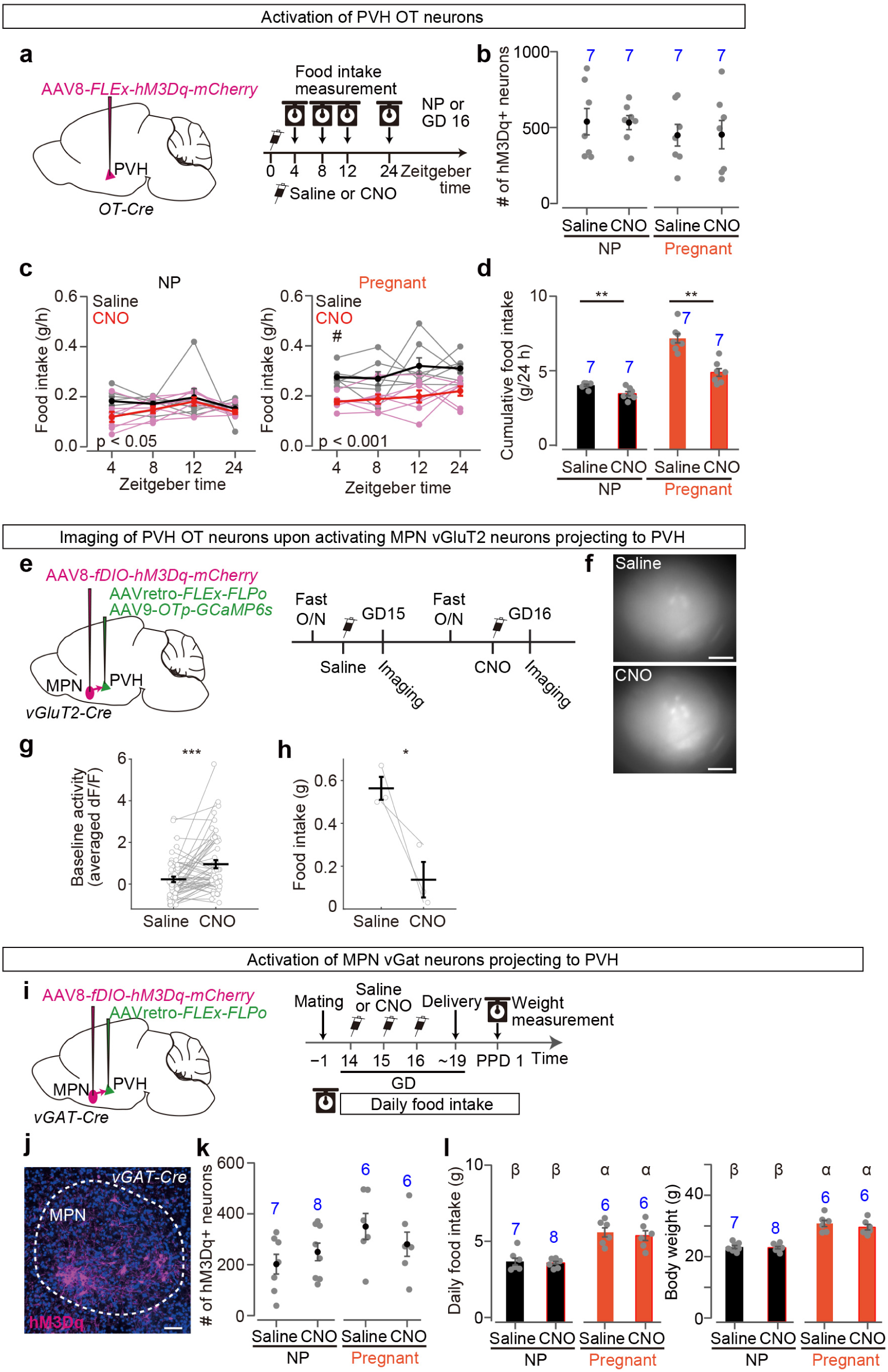
Chemogenetic activation of PVH OT neurons and PVH-projecting *vGAT*+ MPN neurons, related to Fig. 4. **a**, Schematic of the virus injections (left) and the timeline of the experiment (right). Saline or CNO was injected into NP and GD 15 females at Zeitgeber time 0; food intake was measured at 4, 8, 12, and 24 h post-injection. **b**, The number of neurons expressing hM3Dq in the PVH. No significant difference was found among groups by one-way ANOVA. **c**, Hourly food intake (g/h) recorded at 4, 8, 12, and 24 h after injection. #, saline versus CNO, p < 0.05 by significant two-way ANOVA followed by *post-hoc* two-sided Mann-Whitney *U*-test with Bonferroni correction. **d**, Cumulative 24-h food intake (g) was significantly reduced by CNO administration in both NP and GD 15 groups, with a greater effect observed in the GD 15 group. **p < 0.01 by two-sided Mann-Whitney *U*-test. **e**, Schematic of the virus injections and the experimental timeline for chemogenetic manipulation. **f**, Representative images showing baseline Ca^2+^ activity 15 min after saline (left) or CNO (right) injection. **g**, Quantification of baseline activity levels. **h**, Food intake during the imaging session. *p < 0.05 by Welch’s *t*-test. N = 3 mice. **i**, Schematic of the virus injection (left) and the timeline of the experiment (right) for chemogenetic activation of PVH-projecting MPN *vGAT*+ neurons. Saline or CNO was injected over 3 consecutive days from GD 14 to 16 in the pregnant group and the corresponding days in the NP control group. **j**, Representative coronal section of the MPN from a *vGAT-Cre* female expressing hM3Dq (magenta). Blue, DAPI. Scale bar, 30 μm. **k**, Number of neurons expressing hM3Dq in the MPN. **l**, Daily food intake (left) and body weight (right) following chemogenetic activation of PVH-projecting MPN *vGAT*+ neurons. Different letters (α, β) in the upper part of the graph denote significant differences at p < 0.05 by one-way ANOVA with post-hoc Tukey HSD. No significant effect was found by the administration of CNO. Error bars, SEM. The number of animals (N) is indicated in blue.

**Extended Data Fig. 7.**
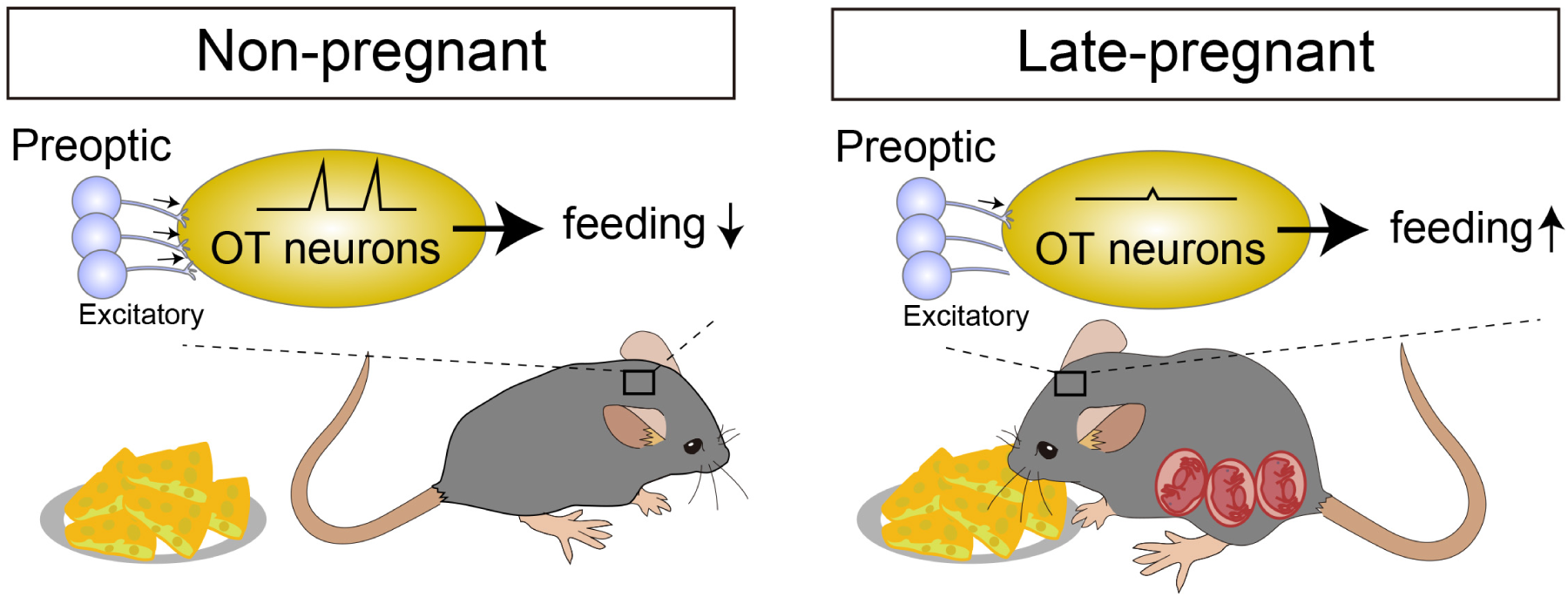
Graphical abstract, related to Figs. 1–4. This study revealed pregnancy-associated neural circuit remodeling upstream of PVH oxytocin (OT) neurons. Specifically, we identified a reduction in excitatory synaptic input from the medial preoptic area to the PVH OT neurons during late pregnancy. This synaptic reorganization is associated with the decreased activity of PVH OT neurons in response to feeding, an anorexigenic function of this population, thereby facilitating increased food intake to meet the metabolic demands of pregnancy.

**Movie S1. Video showing activity of individual PVH OT neurons during feeding by microendoscopy, related to Fig. 3**.

Microendoscopic GCaMP6s-based Ca^2+^ imaging of the PVH OT neurons was performed in non-pregnant female mice. Time zero corresponds to the food supply. Soon after the food was supplied, the animal started feeding, and the PVH OT neurons exhibited Ca^2+^ responses. The movie was presented at 5× real-time speed.

Table S1 List of abbreviations, related to Fig. 1

Table S2 Data set of convergence index, related to Fig. 1

Table S3 Data set of the time course of convergence index, related to Fig. 1

## Notes

### Competing Interest Statement

The authors have declared no competing interest.

